# Longitudinal Changes in Cortical Response Dynamics with Deep Brain Stimulation to the Subcallosal Cingulate for Treatment-Resistant Depression

**DOI:** 10.1101/2025.11.26.690543

**Authors:** Aashna Desai, Tine Van Bogaert, Ki Sueng Choi, Ha Neul Song, Jungho Cha, Sankaraleengam Alagapan, Elisa Xu, Jacob Dahill-Fuchel, Ashan Veerakumar, Brian H. Kopell, Martijn Figee, Patricio Riva-Posse, Chris J. Rozell, Helen Mayberg, Allison C. Waters

**Affiliations:** Nash Family Center for Advanced Circuit Therapeutics, Icahn School of Medicine at Mount Sinai, New York, NY, USA; Department of Psychiatry, Icahn School of Medicine at Mount Sinai, New York, NY, USA; Department of Neurosurgery, Icahn School of Medicine at Mount Sinai, New York, NY, USA; Department of Radiology, Icahn School of Medicine at Mount Sinai, New York, NY, USA; School of Electrical and Computer Engineering, Georgia Institute of Technology, Atlanta, GA, USA; Department of Psychiatry and Behavioral Sciences, Emory University, Atlanta GA, USA; Department of Neuroscience, Icahn School of Medicine at Mount Sinai, New York, NY, USA; Department of Neurology, Icahn School of Medicine at Mount Sinai, New York, NY, USA

**Keywords:** Deep Brain Stimulation, Treatment Resistant Depression, SCC, Tractography, Biomarker, Evoked Potentials

## Abstract

Deep brain stimulation (DBS) of the subcallosal cingulate cortex (SCC) is a promising intervention for treatment-resistant depression, yet objective biomarkers that track recovery remain limited. This study examined longitudinal changes in stimulation-evoked potentials (SEPs) to characterize how SCC-driven cortical communication evolves during treatment. Ten patients across three SCC-DBS trials underwent high-density EEG recordings at 4 and 24 weeks. SEP features were extracted from source-localized SCC signals and related to clinical outcomes and fractional anisotropy (FA) of midcingulate cingulum. Cortical responses showed a consistent reduction in latency and an increase in magnitude over time, indicating faster and stronger electrocortical signaling with chronic stimulation. Higher baseline midcingulate FA predicted greater latency acceleration, linking SEP timing to white-matter integrity. These findings identify temporal SEP dynamics as candidate mechanistic biomarkers that reflect circuit engagement during SCC-DBS and offer a pathway toward physiology-guided optimization of neuromodulation for depression.

**Highlights:**

- Reduced latency of stimulation evoked potential after 6 months of DBS
- Larger cortical evoked response to single pulse perturbation after 6 months of DBS
- Baseline myelin integrity in MCC predicts magnitude of change in EP latency

## Introduction

Treatment resistant depression (TRD) is a neuropsychiatric illness where patients fail to achieve remission despite multiple pharmacologic and psychotherapeutic interventions (1). For these severe, refractory cases, patients are often ECT resistant and deep brain stimulation (DBS) of the subcallosal cingulate cortex (SCC) has emerged as a promising therapeutic strategy (2, 3 4, 5, 6, 7). While SCC-DBS has demonstrated sustained antidepressant benefit and long-term efficacy in a subset of patients and subjective reporting of alleviated symptoms (7), the electrophysiological changes that accompany recovery over the course of treatment remain to be fully characterized. This variability underscores the need for objective neurophysiological biomarkers that can elucidate mechanisms of recovery and guide patient-specific optimization.

Electrophysiological measures, such as stimulation-evoked potentials (SEPs), quantify neural responsiveness and circuit-level engagement (8). SEPs represent time-locked cortical potentials that may reflect in part how stimulation propagates through connected white matter pathways. Waters et al. demonstrated that SEPs elicited by SCC stimulation are reliable and reproducible across sessions, establishing them as a potential biomarker of circuit engagement (9). However, SEP characteristics have not been systematically examined longitudinally to assess how they evolve with clinical recovery or how they relate to the efficiency of communication within and across networks engaged by DBS.

Structural and functional imaging studies have demonstrated that effective SCC stimulation engages distributed fronto-limbic pathways, including projections via the cingulum bundle, uncinate fasciculus, and forceps minor (10). Connectomic targeting approaches have further identified that engagement of these specific white matter bundles is associated with favorable clinical outcomes (11). A study in a non-human primate model showed that chronic SCC-DBS induces selective increases in fractional anisotropy within the midcingulate segment of the cingulum bundle, indicating stimulation-driven remodeling of white matter in downstream tracts (12). This implies an emerging mechanistic theory by which SCC DBS enhances neuronal communication through specific fiber bundles through myelination.

In this study, we utilize deep brain stimulation evoked potentials to measure changes in electrocortical signaling dynamics after six months of treatment for depression with SCC DBS. Specifically, we assess for increase in evoked potential magnitude and latency between two treatment timepoints. We then evaluate these electrophysiological changes as correlates of clinical outcome and fractional anisotropy, a measure of myelin health. We speculate that the speed of stimulation-evoked signal propagation is a function of the brain’s underlying structural connectivity. By linking stimulation-locked neural dynamics with both circuit-level organization and clinical state, this work aims to identify electrophysiological markers that reflect mechanisms of recovery and may ultimately serve as mechanistic biomarkers for DBS efficacy in TRD.

## Methods

### Participants

Ten patients diagnosed with TRD participated in a clinical trial that involved bilateral DBS implantation in the SCC. Four patients were recruited and treated at Emory University School of Medicine, and six additional patients were enrolled at the Icahn School of Medicine at Mount Sinai. Inclusion and exclusion for the Emory cohort are described in Riva-Posse et al. (2018) (11). The corresponding criteria for the Mount Sinai cohort are detailed in Alagappan et al. (2023) (13).

All participants provided written informed consent prior to participation. The study protocol adhered to the ethical standards set forth by the Declaration of Helsinki and received approval from the Institutional Review Boards of Emory University and Mount Sinai. The trial was also approved by the U.S. Food and Drug Administration under a physician-sponsored Investigational Device Exemption (IDE G130107) and registered on ClinicalTrials.gov (NCT01984710).

Participant safety was overseen by the Emory University Department of Psychiatry and Behavioral Sciences Data and Safety Monitoring Board.

**Table 1:** Demographic and Clinical Characteristics of Participants.

| <u>Patient</u> | <u>Sex</u> | <u>Age</u> | <u>Change in HDRS-17 (Baseline to 24 Weeks)</u> |
| --- | --- | --- | --- |
| 1 | F | 65-80 | -12.74 |
| 2 | F | 35-50 | -12 |
| 3 | F | 50-65 | -10.75 |
| 4 | M | 20-35 | -13.5 |
| 5 | M | 50-65 | -19 |
| 6 | F | 35-50 | -18 |
| 7 | F | 50-65 | -13.25 |
| 8 | M | 35-50 | -18.75 |
| 9 | F | 35-40 | -20.75 |
| 10 | M | 20-35 | -15.75 |

HDRS-17: Hamilton Depression Rating Scale

### Tractography-Guided Surgical Implantation of DBS

Each patient underwent surgical implantation of bilateral electrode leads using a tractography-guided approach. This connectome-based targeting approach places the lead at the intersection of three white matter bundles: the cingulum bundle, forceps minor, and the uncinate fasciculus. Bilateral leads (Medtronic, model 3387, spacing 1.5 mm) were implanted and connected to an Activa PC+S in the Emory Cohort and an Activa RC+S in the Sinai Cohort. Postoperative was registered to preoperative CT to confirm lead position relative to the tractography.

### Study Procedure

Patients took part in baseline (presurgical) magnetic resonance imaging, monthly EEG recording sessions and symptom severity measures at each time point. Data from two sessions were utilized in the present analysis. The first session occurred immediately before the activation of therapeutic DBS (about four weeks post-surgery) and the second followed 24 weeks of treatment.

In each EEG session, therapeutic DBS was first turned off in both hemispheres. Participants were instructed to relax the muscles in the shoulders, neck, and face, and to blink, naturally. Unilateral stimulation was delivered to the SCC at 2 Hz in a monopolar configuration with the IPG acting as the anode. The device was voltage-controlled at 6 V in four patients and current-controlled at 6 mA in six patients. Pulse width was set at 90 microseconds. Each contact was stimulated continuously for three minutes, yielding roughly 360 pulses per case. Stimulation cases were tested in ascending order of contact number. After this unipolar stimulation block, therapeutic (clinically prescribed) DBS settings were restored. Data obtained when stimulating on the therapeutic contact of the left hemisphere was retained for the present analysis.

### Symptom Severity

Depression symptom severity was assessed at each time point by a trained clinician using the Hamilton Depression Rating Scale (HDRS-17) which is a clinical questionnaire to assess symptom severity of a patient’s depression (14). Baseline HDRS was calculated as the average score across four weeks preceding surgical implantation.

### Electroencephalography

EEG data were collected using an Electrical Geodesics Inc. system (MagStim-EGI; Eugene, OR, USA), incorporating a 265-channel HydroCel Geodesic Sensor Net together with a Net Amps 400 amplifier. Signals were sampled at 1000 Hz, and recordings were referenced to the vertex reference. Electrode impedances were maintained below 50 kΩ throughout.

### Stimulation Evoked Potentials

Stimulation peak indexing was performed to accurately segment EEG data into time-locked trials corresponding to each DBS pulse. Following segmentation, data were band-pass filtered to remove slow drifts and high-frequency noise (e.g., 1–70 Hz), and bad channels which were identified through manual inspection or automated algorithms detecting signal dropout, excessive amplitude, or flat-lining were excluded from further analysis. Artifact rejection was applied at the trial level to eliminate segments contaminated by non-neural artifacts such as muscle movement, eye blinks, or electrical interference. Evoked responses were averaged across more than 300 trials, resulting in a high signal-to-noise ratio and robust estimation of the stimulation-locked cortical response. Baseline correction was applied to each averaged trial using a pre-stimulation window to ensure that subsequent amplitude measures reflected true post-stimulation deviations in cortical excitability.

### Neural Source Analysis

Neural source analysis was performed to estimate the underlying cortical sources of the stimulation-evoked potentials. Source modeling was carried out using Geosource software (MaStim-EGI, Eugene, OR). sLORETA (standardized low-resolution brain electromagnetic tomography) was selected to solve the inverse problem using a Tikhonov regularization constant of 1 × 10^−2^. Based on the results of Waters et al., (2018), a subset of the 2447 cortical parcellation atlas was selected to represent the SCC region. The root-mean-square average of these dipoles was then exported as a time trace extending 200 ms beyond the stimulation pulse.

### Feature Extraction

To characterize changes in evoked potential amplitude and latency, analyses focused on the most consistent component, which peaked at approximately 40 ms in all patients. Latency was defined relative to component onset, the first trough initiating a rise to component maximum. This measure provides insight into the temporal dynamics of the SCC electrocortical response to a single pulse of DBS. Amplitude of the evoked response was operationalized as the difference between the peak amplitude and the originating trough preceding the peak. This approach allowed us to measure the magnitude of the SCC neural response to single pulses of DBS.

### Fractional Anisotropy

Diffusion-weighted images (DWIs) were acquired on a 3 T Siemens Tim Trio or Prisma scanner using a single-shot spin-echo echo-planar imaging sequence (64 non-collinear directions; five b0 images; b = 1,000 s/mm^2^; 64 axial slices; voxel size = 2 × 2 × 2 mm^3^; TR = 11,300 ms; TE = 90 ms). An additional reverse–phase encoding dataset was collected to correct susceptibility-induced distortions.

All diffusion data were preprocessed using FSL (v6.0). Images were corrected for susceptibility distortion and head motion using *topup* and *eddy*, followed by local tensor fitting to generate voxelwise fractional anisotropy (FA) maps. Individual FA maps were then nonlinearly registered to the FMRIB58 FA template and entered into a standard Tract-Based Spatial Statistics (TBSS) pipeline. A mean FA image was created and thinned to generate a white matter skeleton (threshold FA > 0.2), representing major white matter pathways common to all participants.

For this study, mean FA values were extracted specifically from the mid-cingulate cortex (MCC) region on the FA skeleton. This tract was chosen based on prior macaque SCC-DBS work demonstrating stimulation-induced increases in MCC FA paralleling behavioral recovery (12).

The MCC mask was defined in standard space and intersected with each participant’s skeletonized FA map to obtain subject-specific MCC FA values for group-level analyses (13).

### Statistical Analysis

Analyses were conducted using Python (v3.10). Clinical improvement was evaluated by comparing Hamilton Depression Rating Scale (HDRS) scores at baseline and 24 weeks using a paired *t*-test. For neural data, grand-averaged SCC stimulation-evoked potentials were visualized at 4 and 24 weeks to assess qualitative changes in waveform morphology.

Quantitative comparisons of peak latency and peak magnitude across these two time points were performed using paired *t*-tests (n = 10).

To assess the relationship between white-matter microstructure and SEP changes, we extracted baseline MCC fractional anisotropy (FA) for each participant. For each individual, latency and magnitude differences were computed as the change from 4 weeks to 24 weeks. Linear regression analyses were then performed with baseline FA as the predictor and SEP change (latency or magnitude) as the outcome variables. Pearson correlation coefficients were also calculated to verify the statistical relationships. All tests were two-tailed, with significance set at *p* < 0.05.

## Results

### Longitudinal changes in SCC-evoked potentials across treatment

Using the SCC source montage, we extracted stimulation-evoked potentials (SEPs) from all recording sessions and compared features across early (4-week) and later (24-week) treatment time points.

Grand-average waveforms from left-hemisphere stimulation revealed one consistent peak around 40 ms. Visual inspection suggested a systematic temporal and amplitude shift in this peak over treatment.

Quantitatively, latency decreased significantly from 4 to 24 weeks (*t*(9) = −3.01, *p* = .014; Fig. 2b). Peak-to-trough magnitude increased over the same interval (*t*(9) = 2.36, *p* = .044). These findings indicate that SCC stimulation engages faster and larger cortical responses as treatment progresses.

### Behavioral outcomes across the cohort

All ten patients showed clinical improvement over the 24-week treatment period, with significant reductions in HDRS-17 scores from baseline to 24 weeks (*t*(9) = 14.09, *p* = 1.09 × 10^−7^; Fig. 1b).

**Figure 1.**
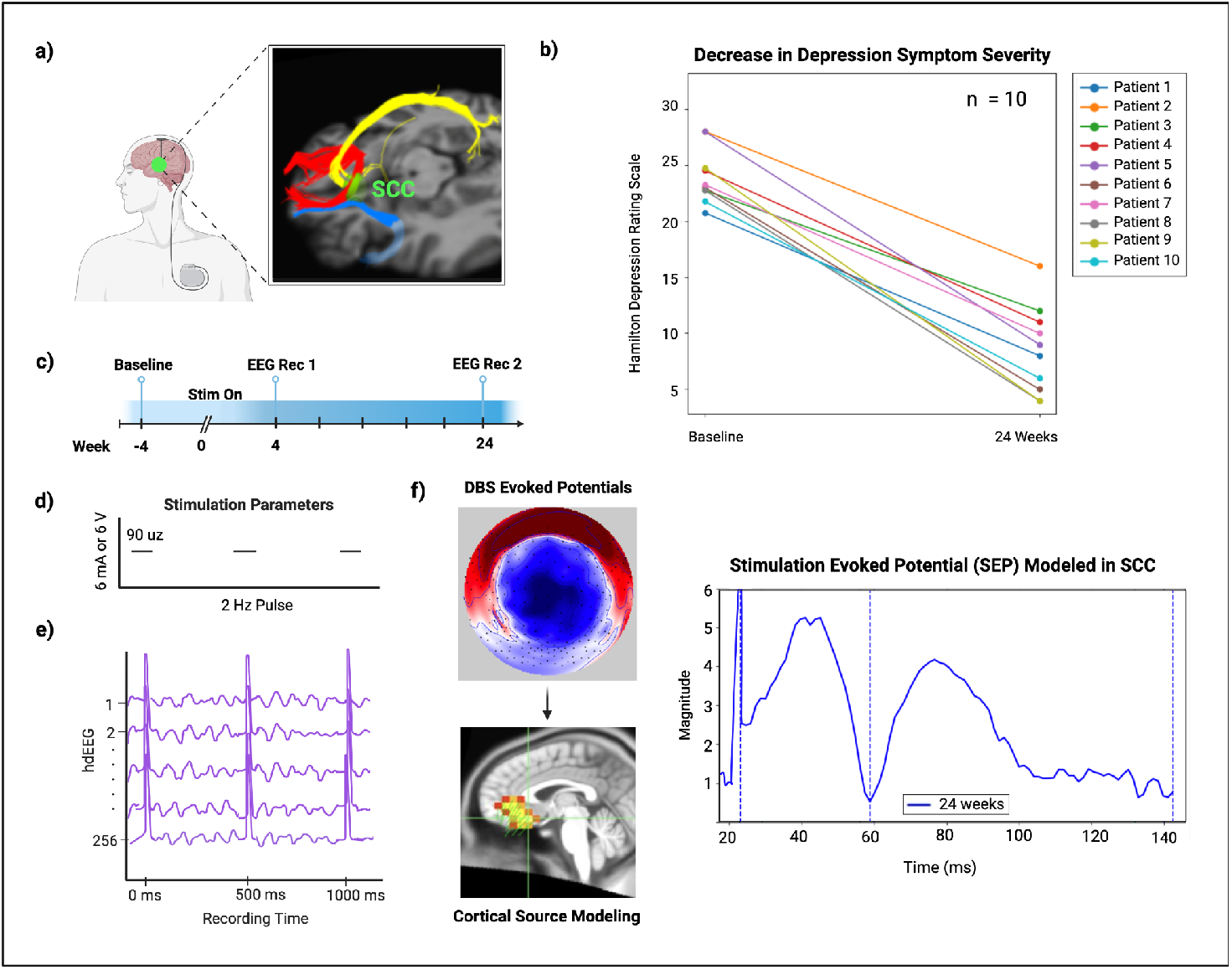
Experimental design and clinical course across the study. (a) Tractography-guided electrode implantation. Bilateral DBS leads were placed at the intersection of the cingulum bundle (yellow), forceps minor (red), and uncinate fasciculus (blue). (b) Longitudinal clinical trajectories showing HDRS-17 scores for all 10 participants from baseline to 24 weeks of stimulation, demonstrating significant clinical improvement across the cohort (*t*(9) = 14.09, *p* = 1.09 × 10^−7^). (c) Study timeline. Participants underwent baseline recordings prior to stimulation onset, followed by electrophysiological recording sessions at 4 and 24 weeks. (d) Stimulation parameters for 2-Hz SCC stimulation (90-μs pulse width; 6 mA or 6 V depending on device). (e) Example of stimulation-evoked responses generated by 2-Hz SCC stimulation. (f) Source-modeling workflow. Whole-brain perturbation maps were reconstructed using sLORETA, and SCC-specific activity was extracted from dipoles approximating the DBS target to derive SEP features.

To test whether SEP features covaried with symptom change, we examined correlations between changes in latency and magnitude with change in HDRS across individuals between 4 weeks and 24 weeks of treatment. Neither relationship reached statistical significance (latency: *r*(8) = −.30, *p* = .328; magnitude: *r*(8) = .38, *p* = .281).

### Relationship between evoked response features and white matter integrity

To test whether efficient SEP propagation reflects the integrity of SCC-connected white-matter pathways, we examined whether baseline fractional anisotropy (FA) of the midcingulate cingulum (MCC) predicted SEP dynamics.

Across participants, higher baseline MCC FA significantly predicted a larger shift toward earlier latency over treatment (*r*(8) = −.68, *p* = .019; Fig. 2c). The relationship between baseline FA and changes in magnitude was not significant (*r*(8) = .19, *p* = .601). These results suggest that the timing of evoked responses may depend on the microstructural strength of midcingulate fibers engaged by SCC stimulation.

**Figure 2:**
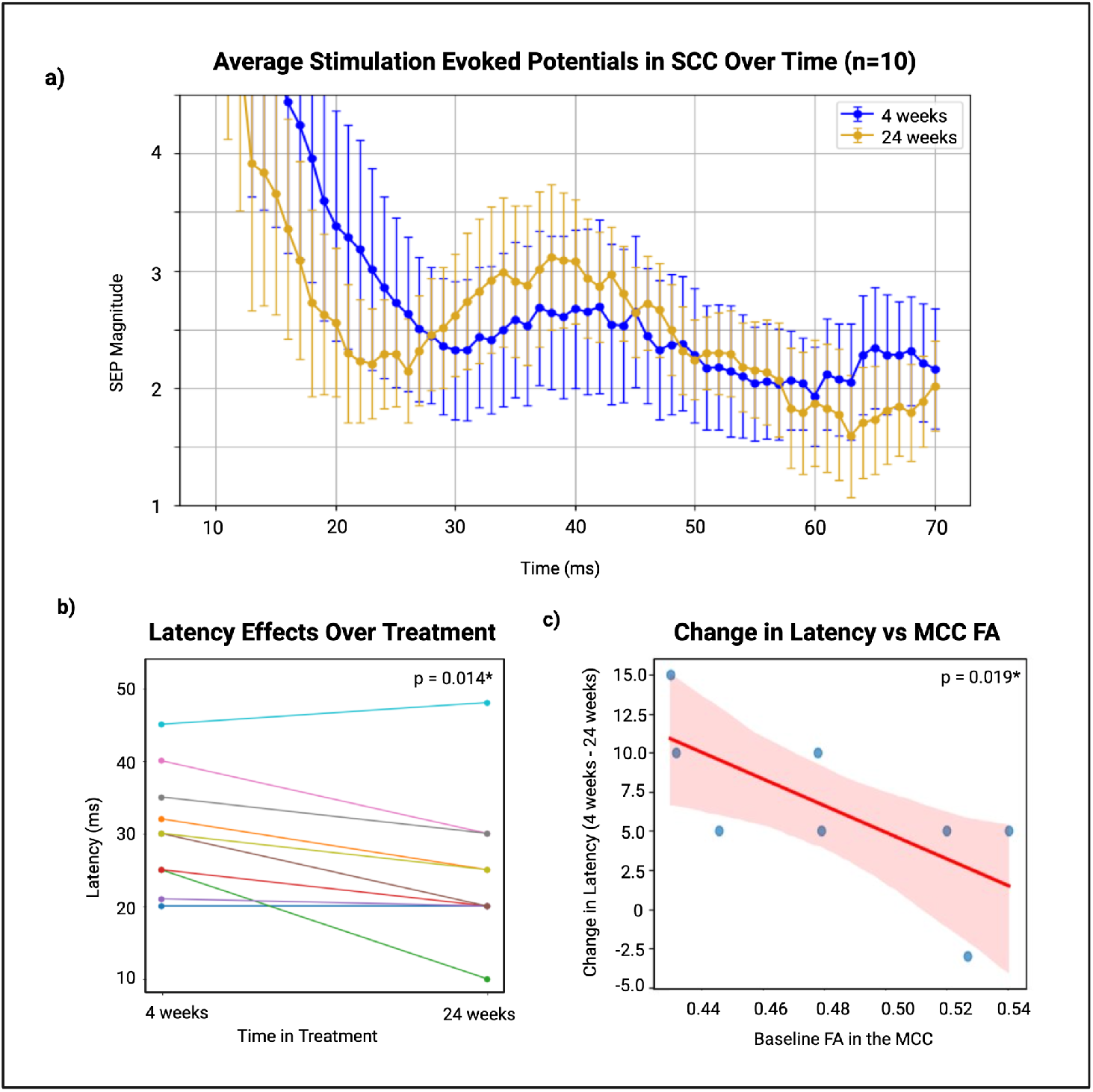
Longitudinal changes in stimulation evoked potentials (SEP) and relationships to white matter integrity. (a) Grand-averaged SEPs from the SCC source following left-hemisphere stimulation at 4 weeks (blue) and 24 weeks (gold), illustrating earlier onset at 24 weeks. Error bars represent the standard error of the mean. (b) Within-participant changes in peak latency from 4 to 24 weeks, showing a significant decrease over time (*t*(9) = −3.01, *p* = .014). (c) Association between baseline fractional anisotropy (FA) in the medial cingulate cortex (MCC) and longitudinal SEP changes. Greater baseline MCC FA was associated with a larger shift toward earlier latency (*r*(8) = −.68, *p* = .019).

## Discussion

This study provides a longitudinal characterization of stimulation-evoked potentials during SCC-DBS for treatment-resistant depression and demonstrates that electrophysiological signatures of SCC network engagement evolve dynamically over the course of treatment. Symptom severity changes confirm therapeutic engagement of SCC-DBS and contextualize the electrophysiological changes under investigation. Across ten participants, we observed a consistent shift toward shorter latency SEPs and an increase in SEP magnitude between 4 and 24 weeks of stimulation. These changes suggest progressive enhancement in the speed and strength of SCC-driven cortical communication as patients move toward clinical recovery.

### Temporal acceleration of evoked responses as a candidate biomarker

Latency shortening was the most robust electrophysiological finding across the cohort. Earlier onset may reflect improved conduction efficiency within circuits engaged by SCC stimulation—potentially indexing neuroplastic processes driven by chronic DBS (12). Prior work has shown that SEPs are reproducible within individuals and sensitive to circuit-level engagement (9); our study extends this by demonstrating that SEP timing changes meaningfully over the course of clinical treatment.

### Relationship between white-matter structure and treatment-dependent SEP changes

A second key finding is that baseline MCC FA predicted the degree of latency shift over treatment. The MCC portion of the cingulum has been identified as a critical downstream tract in SCC-DBS network models (12, 13). Preclinical macaque data show FA increases in the MCC following several months of SCC stimulation, suggesting activity-dependent remodeling along this pathway (12). Although we did not yet acquire longitudinal diffusion data, our results indicate that even baseline MCC integrity partially constrains how efficiently stimulation-evoked signals propagate as treatment progresses. This cross-species correspondence strengthens the interpretation of the latency shift as a tract-dependent circuit biomarker.

### Amplitude changes may reflect cortical excitability rather than tract efficiency

While magnitude increased over time, magnitude changes did not relate to MCC FA. This suggests that amplitude reflects local cortical responsiveness or excitability rather than white-matter conduction efficiency. Prior work in DBS and TMS-evoked responses similarly differentiates timing and amplitude as distinct physiological processes (9, 12, 15, 16) reinforcing the interpretation that latency may serve as the more tract-specific biomarker.

### Implications and future directions

These findings support the use of SEPs as mechanistic biomarkers of SCC-DBS engagement and potentially recovery. An objective neural signal that progresses systematically with treatment could guide parameter optimization and reduce the prolonged trial-and-error characteristic of psychiatric DBS therapy.

Future work should incorporate longitudinal diffusion imaging to test whether electrophysiological acceleration parallels white-matter restructuring in humans, as observed in non-human primates. Additional timepoints—particularly between 4 and 24 weeks—will help map the trajectory of SEP changes and determine whether improvements plateau, continue, or fluctuate with clinical symptoms.

## Conclusion

Together, these results demonstrate that SCC stimulation elicits increasingly rapid and robust cortical responses over the course of treatment, and that the temporal dynamics of these responses relate to the baseline integrity of a key cingulum pathway. These electrophysiological features represent promising candidate biomarkers to quantify and track SCC network engagement during DBS for treatment-resistant depression.

## Acknowledgements

We are deeply grateful to the patients and their families for this commitment and collaboration which made this study possible. We made use of AI technology to improve the readability of the manuscript but not to generate content. We also thank all clinical and research team members whose efforts supported data collection and study coordination. This research was supported by the Brain Research through Advancing Innovation Neurotechnologies (BRAIN) Initiative (UH3NS103550, UH3NS141080), the Hope for Depression Research Foundation and the National Institute of Mental Health (R01MH102238). This content is solely the responsibility of the authors and does not represent the official views of the funding resource parties.

